# Demographic expansions and the emergence of host specialization in genetically distinct strains of the tick-transmitted bacterium *Anaplasma phagocytophilum*

**DOI:** 10.1101/2022.04.05.487247

**Authors:** Matthew L. Aardema, Nina V. Bates, Qiana E. Archer, Friederike D. von Loewenich

**Affiliations:** Department of Biology, Montclair State University, Montclair, NJ, USA; Sackler Institute for Comparative Genomics, American Museum of Natural History, New York, NY, USA; Institute of Virology, University of Mainz, Mainz, Germany

**Keywords:** Host-range, enzootic cycles, arthropod vector, *Capreolus capreolus*, Rickettsiales, Anaplasmataceae

## Abstract

Bacteria species that must obligately replicate in vertebrate host cells make up a large proportion of the prokaryotic pathogens with human and veterinary health implications. In such bacterial taxa, extrinsic processes play an important role in influencing the phylogenetic diversity of viable hosts (‘host range’). These processes include both changes in host population densities and shifts in host geographic distributions. In Europe, distinct genetic strains of the tick-vectored bacterium *Anaplasma phagocytophilum* circulate among mammals in three discrete enzootic cycles. To date, the factors that contributed to the emergence of these strains have been poorly studied. Here we show that the strain which predominately infects roe deer (*Capreolus capreolus*) is evolutionarily derived. Its divergence from a likely host-generalist ancestor occurred after the last glacial maximum as mammal populations, including roe deer, recolonized the European mainland from southern refugia. We also provide evidence that this host-specialist strain’s effective population size (N_e_) has tracked changes in the population of its roe deer host. Specifically, both host and bacterium appear to have undergone substantial increases in N_e_ over the past 1,500 years. In contrast, we show that while it appears to have undergone a major population expansion starting ∼3,500 years ago, in the past 500 years the contemporary host-generalist strain has experienced a substantial reduction in genetic diversity levels, possible as the result of reduced transmission opportunities between competent hosts.

**IMPORTANCE:** The findings of this study are some of the first to examine specific events in the evolution of host specialization in a naturally occurring, obligately intracellular bacterial species. They show that host range shifts and the emergence of host specialization may occur during periods of population growth in a host-generalist ancestor. The results discussed here also show the close correlation between demographic patterns in host and pathogen for a specialist system. These findings have important relevance for our understanding of the evolution of host-range diversity. They may inform future work on host-range dynamics and provide insights for understanding the emergence of pathogens which have human and veterinary health implications.

---

In pathogenic organisms, many distinct extrinsic processes have the potential to alter the phylogenetic extent of viable hosts (‘host range’). These may include changes in the population densities of hosts (Dobson 2004, Woolhouse et al. 2005, Johnson et al. 2020), host geographic range shifts (Thines 2019, Johnson et al. 2020), or the emergence of competitors (Hibbing et al. 2010, Okamoto et al. 2018). However, examples of these processes influencing the host range of specific pathogens remain few (but see Johnson et al. 2020). It is critical to understand the nature of host-range altering processes so as to reduce or mitigate the potential for emergent pathogens to infect and cause illness in humans and domestic animals. As over half of studied viral and bacterial pathogens are host-specialists (Shaw et al. 2020), it is particularly important to examine the processes that contribute to the evolution of restricted host ranges.

The Gram-negative, tick-vectored bacterium *Anaplasma phagocytophilum* is an emerging zoonotic pathogen with human health and veterinary importance (Stuen et al. 2013). In Europe, different strains (or ecotypes) of this obligately intercellular pathogen are genetically distinct, and circulate among mammals in three discrete enzootic cycles (Huhn et al. 2014, Jahfari et al. 2014, Jaarsma et al. 2019, Langenwalder et al. 2020). One of these strains primarily infects nest-living hosts such as voles and shrews (Bown et al. 2009, Huhn et al. 2014, Jahfari et al. 2014, Jaarsma et al. 2019). A second strain is a host-generalist that has been found to infect members of at least five separate mammal orders (Artiodactyla, Eulipotyphla, Perissodactyla, Carnivora, and Primates). Accordingly, *A. phagocytophilum* infections in livestock, companion animals, and humans in Europe are typically caused by this strain (Huhn et al. 2014). The third strain appears to be a host-specialist, being found predominately in roe deer (*Capreolus capreolus*). Despite their substantial difference in host range, the host-generalist and roe deer-specialist strains appear most closely aligned genetically, while the strain found in burrowing mammals is more evolutionary diverged (Huhn et al. 2014, Aardema & von Loewenich 2015).

As a tick-transmitted bacterium, the emergence and maintenance of distinct *A. phagocytophilum* strains could be the result of dynamics in its arthropod vectors. This appears to be a contributing factor in the evolution of the burrowing-mammal strain as its primary vector, *Ixodes trianguliceps*, is found to live almost exclusively in nest and burrows, thus limiting direct transmission potential to larger mammals (Bown et al. 2008, Blaňarová et al. 2014, Mysterud et al. 2015). However, in Europe the other two mammal-infecting strains of *A. phagocytophilum* are both primarily vectored by *I. ricinus*, which has one of the broadest blood-host ranges of any tick species (Estrada-Peña & de la Fuente 2017). While some local population structure and host specialization may occur in *I. ricinus* (McCoy et al. 2013), this is unlikely to contribute to the continued maintenance of the discreet enzootic cycle of the roe deer-specialist strain as it occurs throughout Europe (Huhn et al. 2014, Jahfari et al. 2014, Chastagner 2014, Jaarsama et al. 2019).

If variation in their primary vector does not account for the host-range differences and genetic distinctiveness of the two *I. ricinus*-transmitted strains of *A. phagocytophilum*, then this suggests associations with their mammalian hosts may be a key contributor to strain evolution in this bacterium (Jahfari et al. 2014). During the last glacial maximum in Europe (∼19,000 - 26,500 years ago) (Clark et al. 2009), roe deer were geographically restricted to a few southern refugia (Baker & Hoelzel 2014). As the glaciers receded, roe deer populations were able to spread north into Europe, substantially increasing their geographic range (Matosiuk et al. 2014). An increase in the effective population size (N_e_) of the mainland European roe deer population is estimated to have occurred between 3,932 and 7,919 years ago (Baker & Hoelzel 2014), possibly reflecting this range expansion as well as increases in the continent-wide roe deer census population size. Although relationships between estimates of N_e_ and census population sizes have been shown to lack consistent correlation (Pierson et al. 2008), in natural systems sustained increases in N_e_ generally only occur as the result of a corresponding increase in census population size (Ho & Shapiro 2011, Miller et al. 2021). More recently, the reductions of forest cover corresponding to increases in agricultural practices across Europe starting roughly 1,500 years ago led to a dramatic increase in edge habit (Kaplan et al. 2009, Fyfe et al. 2015, Zanon et al. 2018). This habitat fragmentation likely benefitted roe deer populations and could have resulted in an increase in their population density (Saïd & Servanty 2005, Heurich et al. 2015, Lovari et al. 2017), even as these same habitat alternations reduced the populations of other large mammals (Crees et al. 2016).

It is possible that increases in roe deer population density may have facilitated the emergence of the roe deer-specialist *A. phagocytophilum* strain if host populations were sufficiently large enough to facilitate transmission predominately between these ungulates. In this scenario, selection among linages of *A. phagocytophilum* would have favored any that harbored genetic variation that improved infection capabilities in roe deer. Such selection may have ultimately led to reduced ability to infect other mammals, either through antagonistic pleiotropy whereby adaptions for improved roe deer infection came at the cost of infection ability for other species, or else through the accumulation, via genetic drift, of mutations that reduced infection capabilities in other hosts (Cooper & Lenski 2000).

We can examine a scenario of derived *A. phagocytophilum* host specialization in conjunction with changes in European roe deer populations by testing several corresponding hypotheses. The first of these is that the roe deer-specialist strain is evolutionary derived, having evolved from a more generalist strain that likely resembled the contemporary host-generalist *A. phagocytophilum*. The second hypothesis is that this divergence occurred with increases in the roe deer population of Europe, possibly either as it recolonized the continent following the last glacial maximum (∼3,932 - 7,919 years ago) (Baker & Hoelzel 2014), or else during reductions in forest cover brought about by increases in agriculture (starting ∼1,500 years ago) (Kaplan et al. 2009, Fyfe et al. 2015, Zanon et al. 2018). Finally, the third hypothesis is that after the evolution of the roe deer-specialist strain, changes in this strains’ effective population size would be closely correlated with changes in roe deer population sizes. Given the apparent larger N_e_ observed in the contemporary roe deer-specialist strain (Aardema & von Loewenich 2015), evidence in support of this third hypothesis would also suggest that host specialization has allowed this strain to reach higher population densities than the generalist strain. This could be attributed to greater within-host population sizes and/or an increased frequency of transmission between viable hosts (Dobson 2004, Toft and Andersson 2010). Such an observation would support the possibility that host specialization can correlate with increases in overall fitness for a pathogenic bacterial population.

## RESULTS

### Sample Clustering

To test the three hypotheses described above, we leveraged previously published DNA variation found in seven partial gene sequences amplified from 278 *A. phagocytophilum* samples isolated either from a mammalian host or *I. ricinus* vector (Huhn et al. 2014). We initially performed a nonparametric, principal component analysis (PCA) to confirm sample assignment to one of the distinct, mammal-infecting strains previously identified (Huhn et al. 2014). As predicted, our PCA clearly delineated three discrete clusters among the samples assessed (Fig. 1). The first principal component accounted for 37.2% of the genetic variation observed, and separated the burrowing-mammal strain from the two *I. ricinus*-vectored strains. The second principal component accounted for 8.9% of the genetic variation, and separated the roe deer-specialist strain from the host-generalist strain. All samples clearly fell into one of the three clusters, with no ambiguous, intermediate samples. Of the 278 total samples used in this analysis, there were 20 representing the burrowing-mammal strain, 24 representing the roe deer-specialist strain, and 234 representing the host-generalist strain. All subsequent analyses utilized sample strain designations as determined from this PCA.

**FIG 1.**
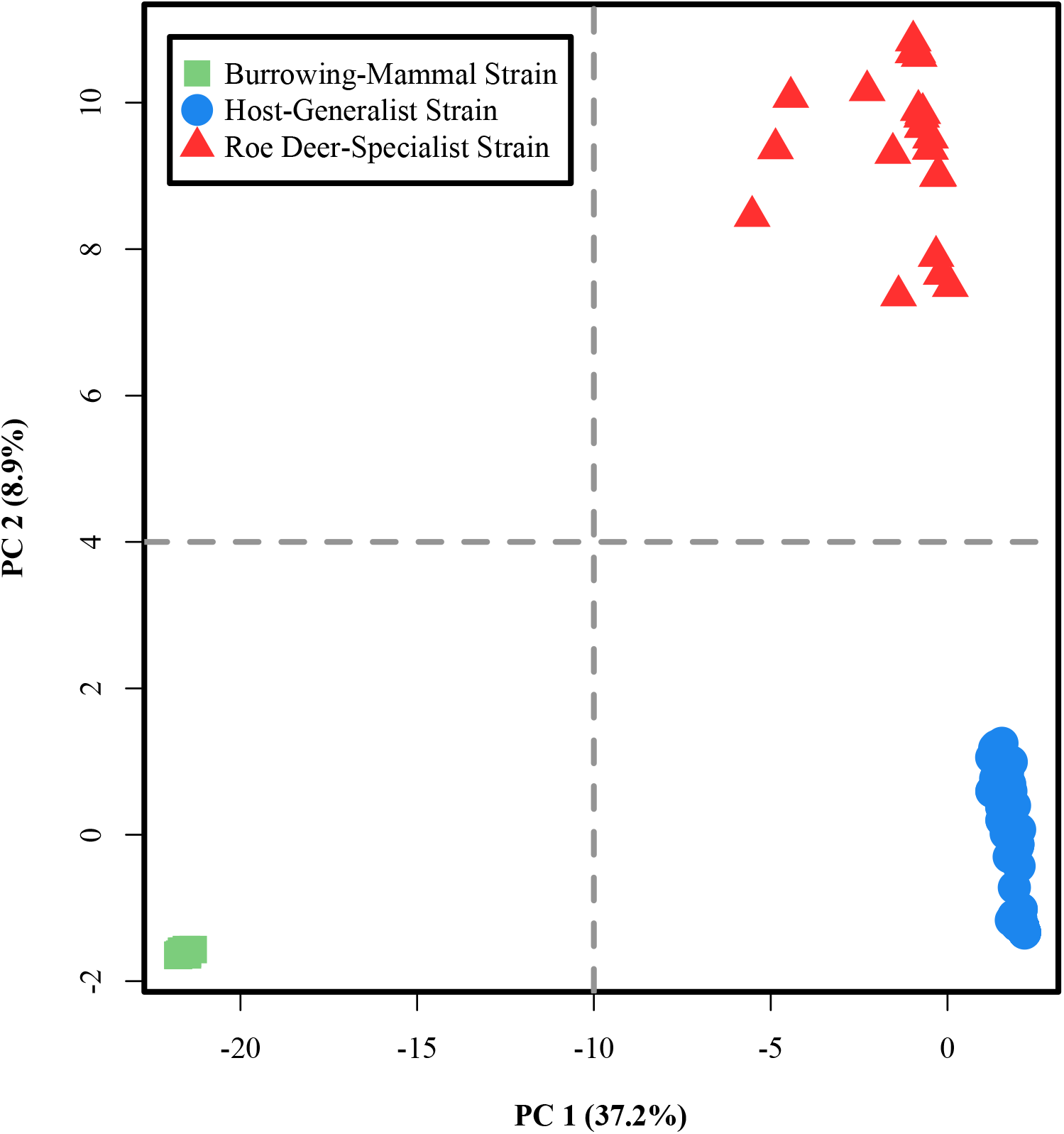
Results from a principal component analysis (PCA) based on concatenated data from seven previously published genetic regions (see text). Shown are the first and second principal components (PC1 and PC2, respectively). This original image was produced using the R package Adegent v. 2.1.3 (Jombart & Ahmed 2011), as implemented in R v. 4.0.2 (R Core Team 2020). Each color/shape combination represents one of the three mammal-infecting strains of *A. phagocytophilum* circulating in Europe.

### Evolutionary Pattern of Strain Divergence

We hypothesized that the contemporary roe deer-specialist strain evolved from an ancestral host-generalist strain, and may accordingly be more derived relative to the contemporary host-generalist strain. When comparing two closely related, contemporary populations, the one which has diverged more from their shared, common ancestor should harbor the signature of this greater divergence in patterns of DNA evolution, specifically the presence of derived alleles (Green et al. 2010, Lachance et al. 2012). During host-range shifts, an excess of derived alleles could emerge and be maintained either because they are directly selectively advantageous in the environment, or because they were physically linked to other adaptive variants (Karlsson et al. 2014, Didelot et al. 2016).

Previous work (Huhn et al. 2014, Jahfari et al. 2014, Aardema & von Loewenich 2015), and our nonparametric analysis (Fig. 1), indicate that the roe deer-specialist and host-generalist strains are most closely related to one another and the burrowing-mammal strain is more evolutionary diverged. We therefore utilized the unique haplotypes observed among samples of the burrowing-mammal strain for outgroup comparisons, assuming that a nucleotide shared by this strain and one or both *I. ricinus*-vectored strains is the ancestral state at this position. Accordingly, an alternative allele in one of the two focal strains is assumed to be derived. Using this conceptual framework, our pair-wise comparisons of each strain’s unique haplotypes showed that the roe deer-specialist strain harbors a greater number of observed, derived changes across the seven examined genetic regions, considering both DNA variation that resulted in an amino acid change (‘replacement’ sites; Fig. 2a), and variation that did not alter the protein sequence (‘silent’ sites; Fig. 2b). For replacement sites, there was an average of 6.53 (SD: 2.24) derived alleles per sample in the roe deer-specialist strain, whereas in the host-generalist strain there was an average of 0.94 (SD: 0.91). These differences were statistically different from one another (t_14079_ = 303.8, P < 0.001). For silent substitutions, the roe deer-specialist samples had an average of 35.54 (SD:11.18) derived alleles, while the host-generalist had an average of 32.22 (SD:13.59) derived alleles. Again, these differences were statistically significant (t_14079_ = 54.6, P < 0.001).

**FIG 2.**
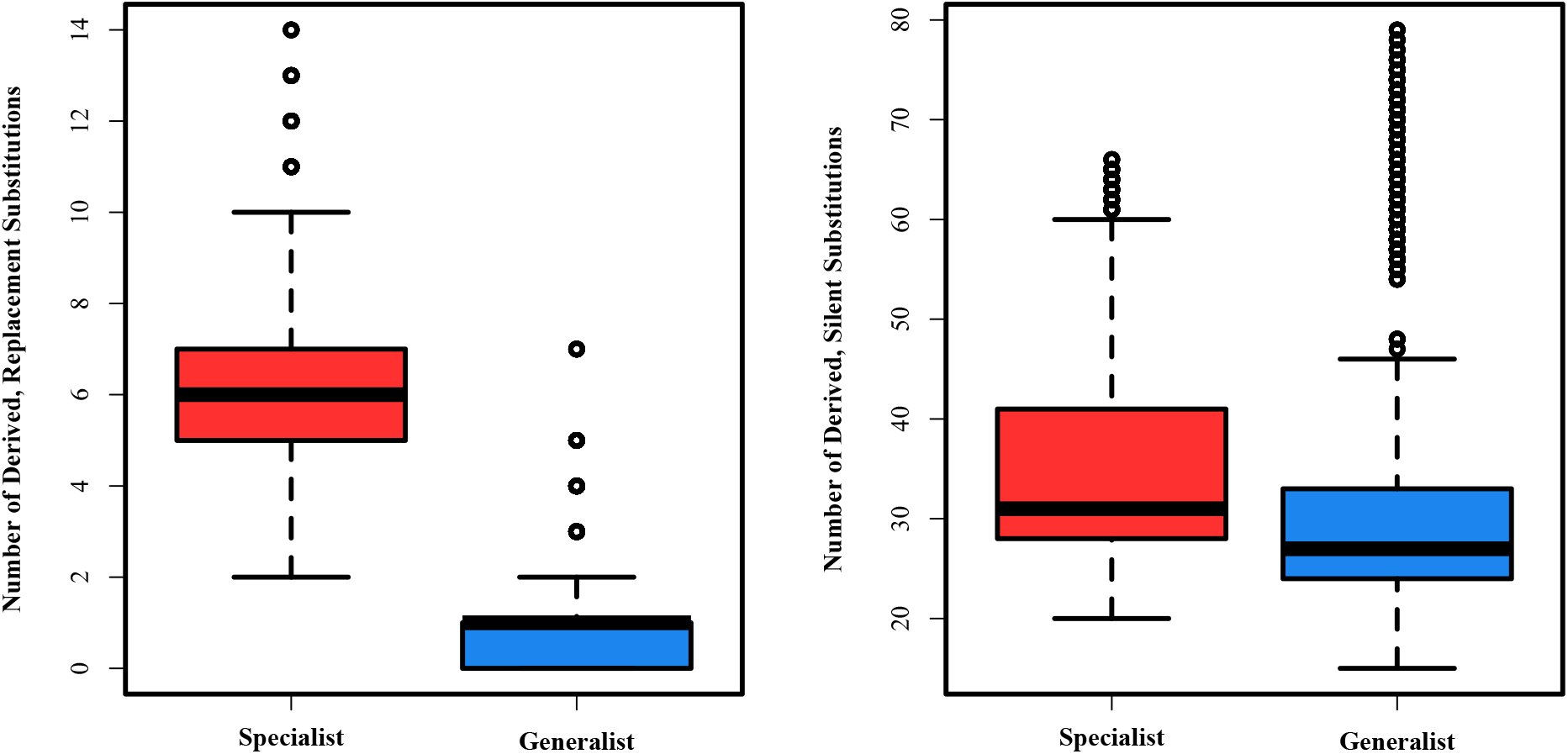
Box and whisker plots indicating the number of observed, derived replacement (a) and silent (b) substitutions observed in all pairwise comparisons of each unique haplotype from the roe deer-specialist and host-generalist strains. Polarization to define the ancestral state was done in comparison to the unique haplotypes of the burrowing-mammal strain. The thick horizontal black bars indicate the observed median number of substitutions. The boxes show the second and third quartiles, while the whickers indicate the first and fourth quartiles. Outlying observations are indicated by the open circles. Circles may represent more than one observation. Both comparisons between the roe deer-specialist strain and the generalist strain were statistically significant (p < 0.001; see text for more details).

### Strain Divergence Times

We estimated approximate splitting times for the three strains, with the goal of assessing how potential changes in European roe deer populations may have correlated with strain divergence in *A. phagocytophilum*. To make these estimates, we used a Bayesian coalescent method, implemented in BEAST, and calibrated to absolute dates based on previously estimated divergence between *A. phagocytophilum* and the outgroup taxon *A. marginale* (Foley et al. 2008, de la Fuente et al. 2015). Our analysis recapitulated the predicted phylogenetic relationships between *A. marginale* and *A*. phagocytophilum, as well as those between the three strains. *A. marginale* was the clear outgroup to all *A. phagocytophilum* haplotypes (Fig. 3a). Within the *A. phagocytophilum* haplotypes, those assigned to the burrowing-mammal strain clustered with one another and collectively were distinct from the *I. ricinus*-vectored strain haplotypes (Fig. 3b). The haplotypes assigned to the host-generalist and roe deer-specialist strains respectively each formed distinct and monophyletic clusters, and were sister to one another. We estimated that the most recent common ancestor of all three strains was present 3,330 years ago, with a range based on the 95% highest posterior density (HPD) from 529 to 8,061 years. Our estimate for the most recent common ancestor of the two *I. ricinus*-vectored strains was 2,970 years ago with a 95% HPD of 454 to 7,240 years.

**FIG 3.**
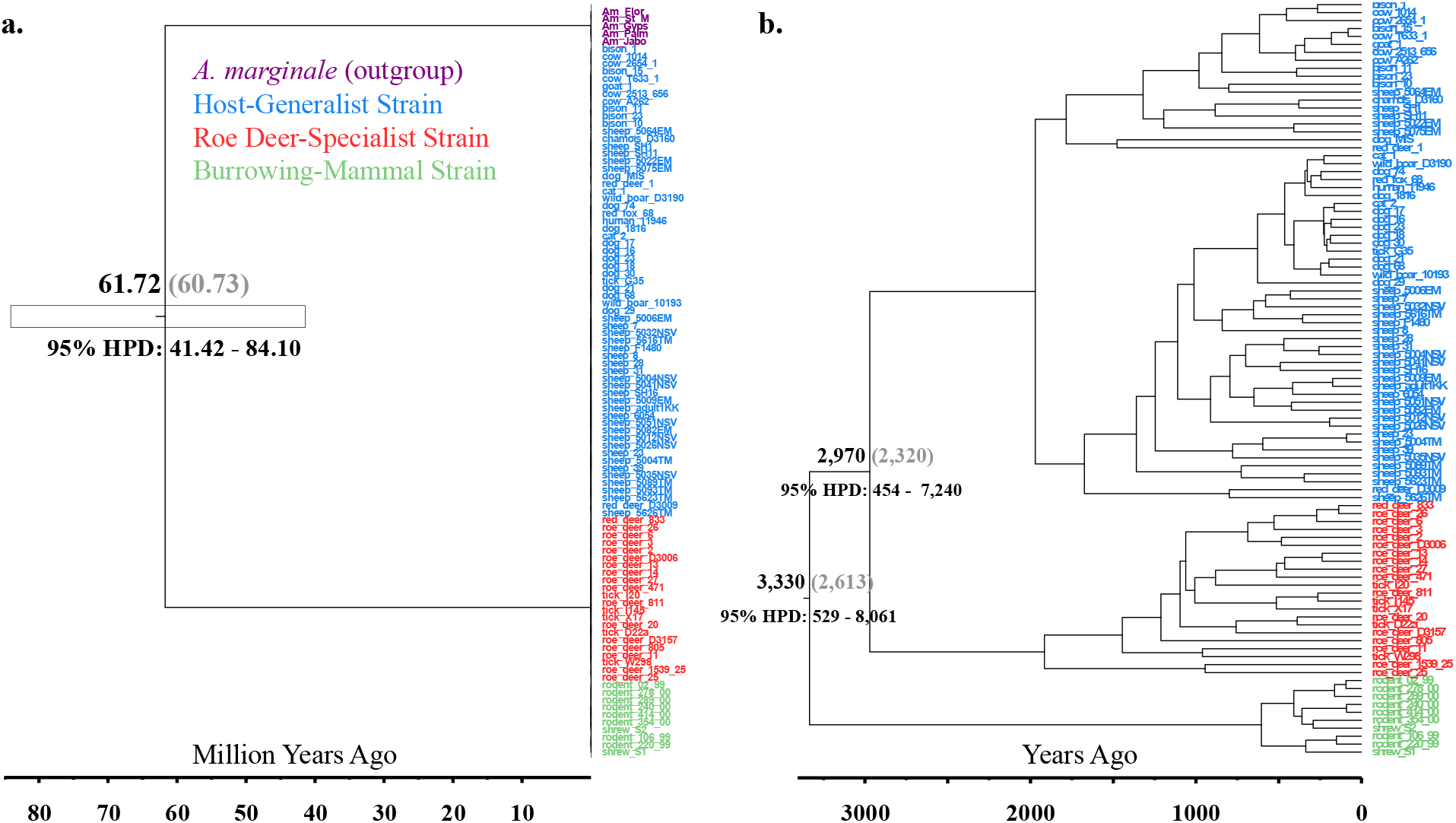
Divergence time estimates for the three strains. a) tree including the outgroup *A. marginale*. b) Subtree with only samples of *A. phagocytophilum*. In both trees the mean estimated divergence time is shown above the node (in black text), followed in parentheses by the median estimated divergence time (in grey text). The 95% HPD is given below the node. Only estimates for nodes representing species/strain divergences are shown. The scale bars at the bottom of each tree indicate time in millions of years (a) or years (b). Duplicate haplotypes were removed prior to analysis.

### Demographic changes in *I. ricinus*-vectored strains and host

Given the relatively recent split observed between the two *I. ricinus-*vectored strains (see above), and the apparent differences in their contemporary effective population sizes (N_e_) (Aardema & von Loewenich 2015), we wanted to determine if these two strains have distinct demographic histories. We also wanted to examine potential correlations between demographic changes in the roe deer-specialist strain and changes in the population density of its mammalian host.

To estimate temporal changes in N_e_ for the two *I. ricinus*-vectored strains, we applied an extended Bayesian skyline model as implemented in BEAST to separate subsets of the data containing only samples from each strain. A clock rate of 8.5 × 10^−8^ changes per generation was applied to the third codon position of our concatenated dataset, and rates for the other two positions were estimated. To convert estimates of change in N_e_ from generations to absolute time, we assumed 100 bacterial generations per year. Our results suggest that the roe deer-specialist strain underwent a substantial population expansion starting approximately 1,150 years ago (Fig 4). The host-generalist strain appears to have a more complex demographic history with an apparent increase in N_e_ starting around 3,500 years ago, followed by a more recent, rapid reduction starting approximately 500 years ago.

**FIG 4.**
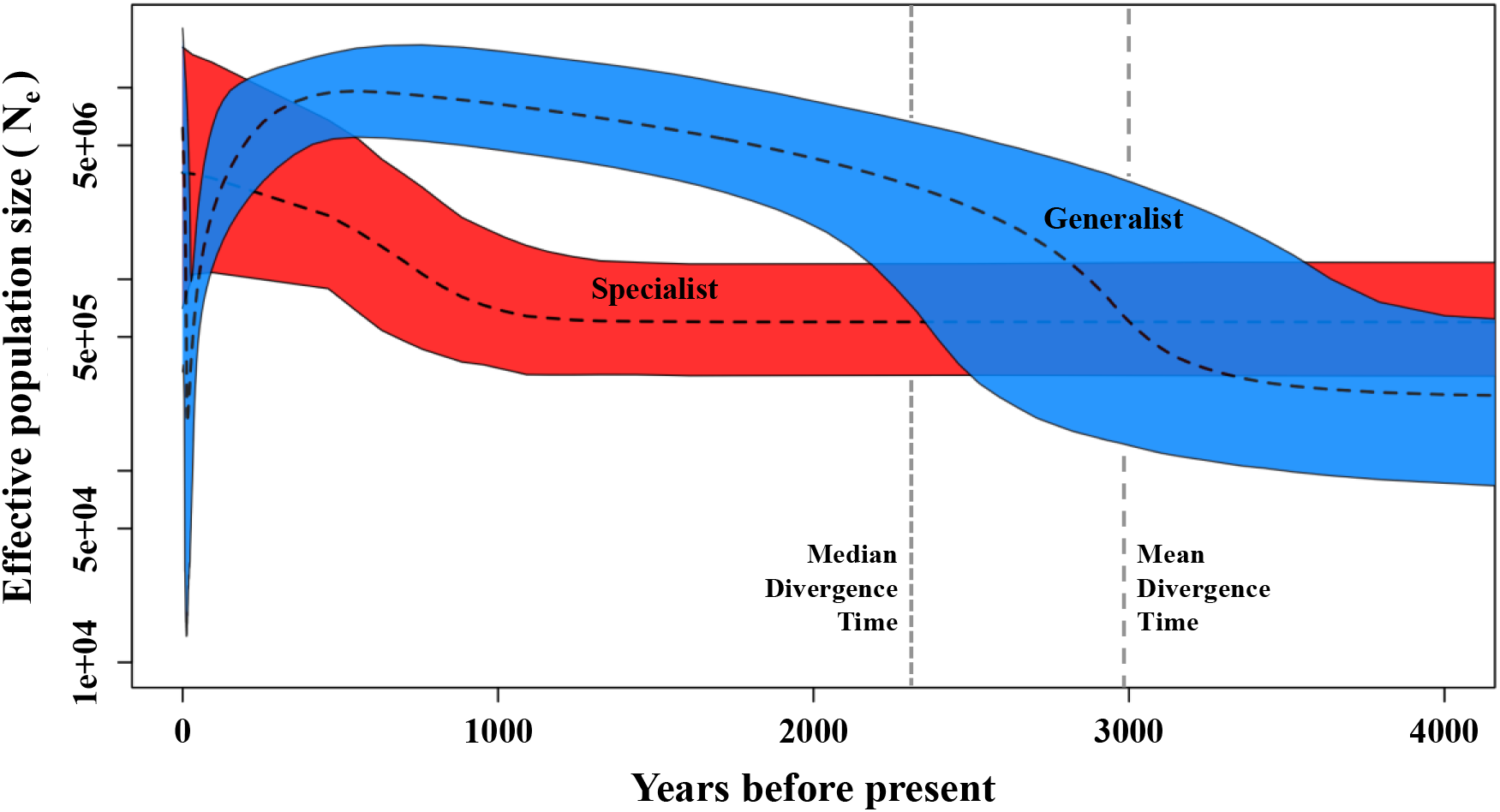
Extended Bayesian skyline plot (EBSP) results for the two *I. ricinus*-vectored strains showing changes in effective population size (N_e_) over time. Median results are shown by the black dashed lines, with the upper and lower 95% highest posterior density (HPD) are indicated by the colored areas contained within the thin solid lines. Changes in the roe deer-specialist strain’s N_e_ are indicated in red, and changes in the host-generalist strain’s N_e_ are indicated in blue. The original timeline was determined in terms of number of generations and this was converted to years assuming 100 generations per year. The timeline has been restricted to 4,000 years. Also shown are the median (finely dashed vertical gray line) and mean (coarsely dashed vertical gray line) divergence estimates from our Bayesian coalescent analysis (see text and Fig. 3 for more details).

We also examined recent demographic changes in the roe deer population of Europe. To do this, we utilized previously published sequences of the mitochondrial cytochrome b (cytb) gene amplified from 46 roe deer from Poland (Matosiuk et al. 2014). It was preferable to use samples from a single country to reduce potential confounding effects of population structure on estimates of demographic history (Heller et al. 2013). Our results indicated a substantial increase in N_e_ for European roe deer starting approximately 1,500 years ago (Fig. 5), with a contemporary N_e_ estimated to be over 20 times as large as that at the start of this apparent demographic expansion.

**FIG 5.**
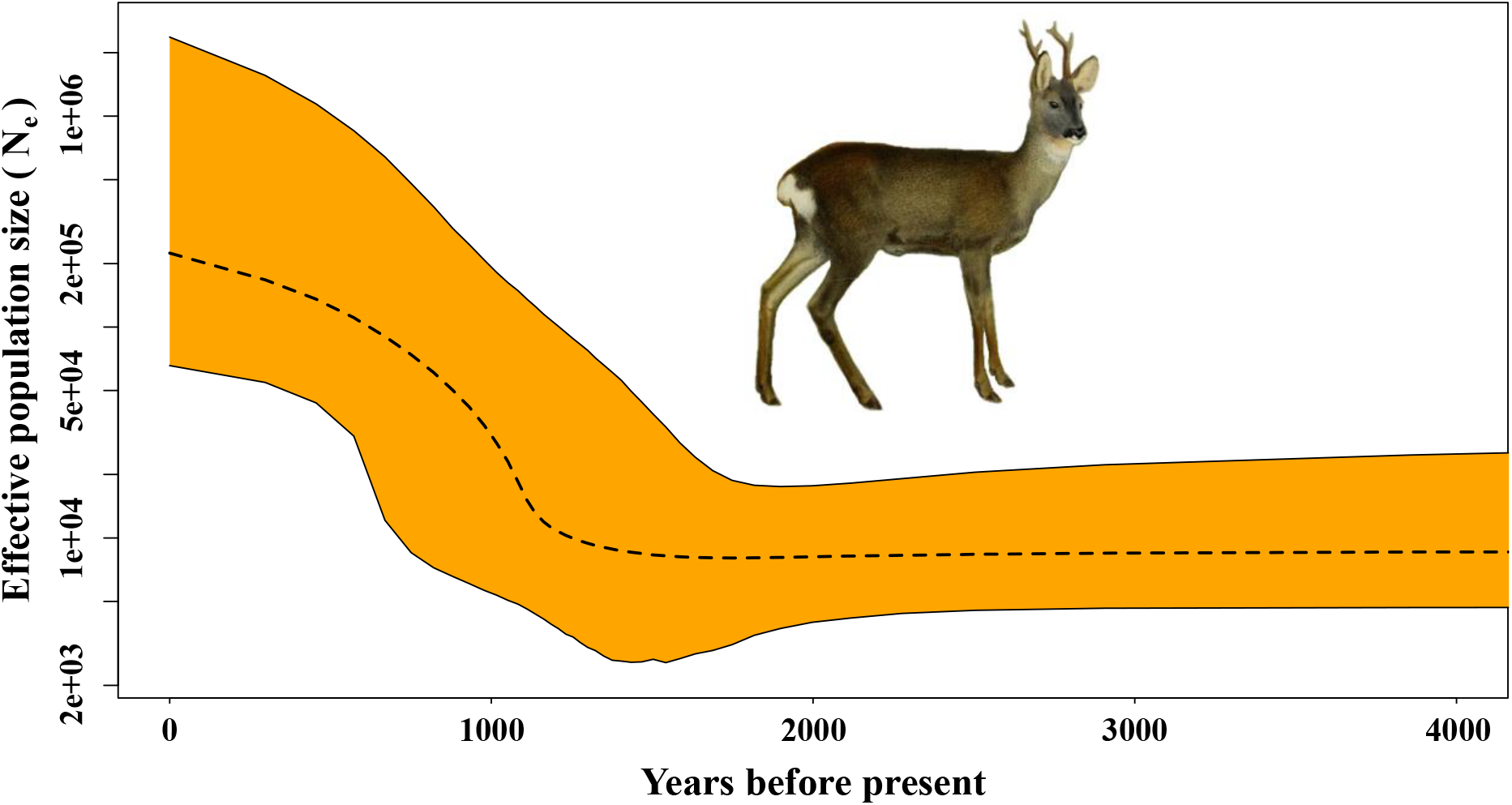
Extended Bayesian skyline plot (EBSP) results for roe deer (*Capreolus capreolus*) showing changes in effective population size (N_e_) over time. Median results are shown by the black dashed line, with the upper and lower 95% highest posterior density (HPD) are indicated by the orange areas contained within the thin solid lines. The timeline has been restricted to 4000 years. Image of a male roe deer by Richard Lydekker, Public domain, via Wikimedia Commons (https://upload.wikimedia.org/wikipedia/commons/b/bd/The_deer_of_all_lands_%281898%29_European_roe_deer_white_background.png).

## DISCUSSION

We postulated that the two contemporary *I. ricinus*-vectored strains of *A. phagocytophilum* share a common ancestor that had a relatively broad host range, and that the roe deer-specialist strain diverged from this ancestral host-generalist. Although we predicted that the contemporary roe deer-specialist strain was evolutionary more derived, it is possible that the contemporary host-generalist strain could have evolved from a more specialized population (Johnson et al. 2009). Nonetheless, our comparison of derived alleles in the host-generalist and roe deer-specialist strains supported our hypothesis concerning host-range evolution, and indicates the specialist strain evolved from a population of *A. phagocytophilum* that likely resembled the contemporary host-generalist with a broad host range (Fig. 2). In turn, this observation suggests that specific changes in the roe deer population of Europe facilitated the emergence and evolution of the roe deer-specialist strain.

One such change might have been the geographic expansion of roe deer in Europe after the last glacial maximum. This recolonization event correlates with a dramatic increase in the N_e_ of mainland European roe deer and is estimated to have occurred between 3,932 and 7,919 years ago (Baker & Hoelzel 2014). It is also possible that an increase in roe deer population density (independent of range expansion) may have contributed to the divergence of the two *I. ricinus*-vectored strains of European *A. phagocytophilum*. Such a density increase likely occurred in conjunction with more intensive agricultural practices in Europe starting around 1,500 years ago (Fyfe et al. 2015, Zanon et al. 2018). Our estimation of the split between the two *I. ricinus*-vectored strains suggests they last shared a common ancestor ∼2,970 years ago, with a range between 454 and 7,240 years (based on the 95% HPD, Fig. 3). This estimate, combined with the earlier conclusion that the roe deer-specialist is the more derived strain, suggests that host specialization followed the geographic range expansion of roe deer throughout Europe after the last glacial maximum. Two *I. ricinus*-vectored populations of *A. phagocytophilum* resembling the contemporary strains had likely already evolved prior to potential increases in roe deer population densities that may have occurred with more recent shifts in the vegetation communities throughout Europe.

While its divergence appears to have occurred earlier, our analysis of patterns of demographic change in the roe deer-specialist strain in relation to changes in the European roe deer population suggest a strong correlation between the two. Specifically, the N_e_ of both appears to have dramatically increased around 1,500 years ago (Figs. 4, 5). These observations support our third hypothesis that changes in the population density of roe deer correlate with an increase in the N_e_ of the roe deer-specialist strain. However, it is possible that these two seemingly congruent changes in N_e_ are coincidental and unrelated. It is also possible that additional factors not considered here were the primary drivers of the effective population size increases in both roe deer and the roe deer-specialist strain of *A. phagocytophilum*, and that the former is not directly the cause of the latter. Nonetheless, if future work does support a causal relationship between host and pathogen in this system, then the apparently strong genetic tracking of its host population by this strain of bacterium strengthens confidence that it is truly a host-specialist, highly adapted to infecting roe deer. It also provides evidence that host specialization can correlate with higher fitness in a pathogenic bacterial population.

Among possible European mammalian hosts, it is perhaps not surprising that roe deer were the species that came to support a specialist strain of *A. phagocytophilum*. Unlike many other mammal species across the European continent, roe deer appear to have benefitted from growth in the human population and the corresponding increased intensity of agricultural (Saïd & Servanty 2005, Heurich et al. 2015, Lovari et al. 2017). Increases in roe deer population density likely meant that *A. phagocytophilum* transmission events became increasingly likely to occur from one roe deer to another. As lineages of *A. phagocytophilum* progressively found themselves only in roe deer, this could have produced strong selection for improved abilities to infect this host, possible at the expense of infection capabilities in other mammals.

The host-generalist strain looks to have experienced a more complicated demographic history (Fig. 4). Intriguingly, it appears to have been in the midst of a substantial population increase when it last shared a common ancestor with the roe deer-specialist strain. This may mean that the emergence of the specialist strain occurred in part because the *A. phagocytophilum* population in Europe at this time was experiencing extensive growth, possibly as the result of increased transmission between hosts. This increased transmission may have been facilitated by range expansions of most mammal species at this time as new habit became available when the glacial ice retreated north (Hewitt 1999). Increased transmission and greater bacterial populations sizes have been shown to correlate with genetic diversity and strain divergence/specialization in pathogenic bacteria (e.g., Ueti et al. 2012, Pollitt et al. 2014, Chavhan et al. 2020). While currently speculative, this scenario would explain both the patterns of strain divergence in European *A. phagocytophilum* and the major increase in the N_e_ of the host-generalist strain during this period.

More contemporarily, the host-generalist strain appears to have undergone a major reduction in N_e_, possibly followed by some very recent recovery (Fig. 4). A reduction in N_e_ could occur if its rate of transmission decreased (Bobay & Ochman 2018). It could also occur in a bacterial population if one or more advantageous alleles became strongly favored and this generated a selective sweep (Bendall et al. 2016). For a bacterial strain with a wide geographic distribution and a broad host range, a selective sweep appears to be the less probable of these two scenarios. Therefore, if we postulate a decrease in the population size of the host-generalist strain, this again points to the importance of host population dynamics in the demography of European *A. phagocytophilum*. While the host-generalist strain can infect humans as well as a wide variety of livestock and companion animal species, its predominant natural reservoirs likely include European bison, wild boars, hedgehogs, and possibly red deer (Huhn et al. 2014, Jahfari et al. 2014, Dugat et al. 2015). Unlike roe deer, many of these species have experienced substantial population declines over the last several centuries in Europe (Scandura et al. 2008, Kuemmerle et al. 2012, Rosvold et al. 2012, Crees et al 2016, Rasmussen et al. 2020).

One caveat of this study that must be discussed is the important differences between effective population sizes (N_e_) and census population sizes. Here we have considered changes in N_e_ modeled by other researchers and carried out our own modeling of changes in N_e_. Effective population sizes are almost always smaller than census population sizes and can decrease for a variety of reasons including genetic drift, selection, or migration (McInerney et al. 2017, Bobay & Ochman 2018). However, with the exception of a decreased N_e_ modeled in the host-generalist strain, the observations of change in N_e_ considered here all suggest increases over time (Figs. 4, 5). Prolonged increases in N_e_ are almost always the result of population growth and therefore should accurately reflect increases in population sizes or rates of transmission (Gillespie 2004, Ho & Shapiro 2011, Hague & Routman 2016, Bobay & Ochman 2014). Nonetheless, it is also possible that the observed increases in *A. phagocytophilum* reflect greater rates of gene flow between populations (Charlesworth 2009, Hague & Routman 2016, Willi et al. 2020).

Another important limitation of this study is the reliance on poorly-substantiated estimates of sequence evolution in bacteria and roe deer to infer patterns of demographic change (Ochman et al. 1999, Nabholz et al. 2009). While the qualitative patterns of change are unlikely to be influenced by these estimates, the timing of such events could shift depending on the mutation rate used. Until more refined and taxon-specific rates become known, this limitation is unavoidable.

One central question that remains is the relative scarcity of the host-generalist strain in roe deer. Although samples found in roe deer have been attributed to the host-generalist strain, such infections are far less common in this host than the specialist strain (Jahfari et al. 2014, Langewalder 2020). Given the phylogenetic diversity of species the host-generalist strain can infect (including other Artiodactyla), it seems likely the host-generalist has the capacity to establish infections in roe deer. This suggests that competition with the roe deer-specialist strain may contribute to the generalist’s absence from this host. Additional work will be required to further elucidate this possibility.

In conclusion, we have shown that among *I. ricinus*-vectored populations of *A. phagocytophilum* in Europe, the roe deer-specialist strain is the more derived. We have also shown that its divergence from a likely ancestral host-generalist occurred after the last glacial maximum as mammal populations (including roe deer) recolonized the European mainland. Finally, we have provided evidence that the strain of *A. phagocytophilum* that infects roe deer in Europe is truly a specialist of this host, and that changes in the population of roe deer strongly correlate with the population dynamics of this bacterium. Comparisons of these closely related strains will help us better understand the ways in which host specialization influences ecology, demography, and genomic evolution in pathogenic bacteria. Because they are so closely related, examination of these two strains should allow assessments to exclude some of the large, species-specific differences that may correlate with host-range evolution such as motility or transmission mode (Shaw et al. 2020). Of particular interest are the processes in the roe deer-specialist that may have led to a reduction of infection capabilities in other hosts (Cooper & Lenski, 2000). During host specialization deleterious mutations may have become fixed by genetic drift in genes important for infection in other hosts but not in roe deer. It is also possible that genetic trade-offs occurred for improved infection capabilities in roe deer at the expenses of infection potential more broadly (i.e., ‘antagonistic pleiotropy’). In this case, specific mutations that improved roe deer infection would be detrimental to infection of other hosts. While it can be difficult to distinguish between these processes, the two *I. ricinus*-transmitted *A. phagocytophilum* strains represent an excellent system in which to examine this problem. Future research may also allow a better understanding of the importance of genetic changes, including possible gene loss, in facilitating host specialization (Parkhill et al. 2003, Cummings et al. 2004). An understanding of the process which occur during host specialization will have important ramifications for understanding broad patterns of pathogen evolution.

## MATERIALS AND METHODS

### Dataset and sample clustering

Throughout this study we utilized partial sequences from seven *A. phagocytophilum* genes (*atpA, dnaN, fumC, glyA, mdh, pheS*, and *sucA*), representing 2,877 nucleotides in total (GenBank accession numbers KF242733-KF245413). These data were sequenced from *A. phagocytophilum* DNA isolated from 17 different mammalian hosts and *Ixodes ricinus* ticks, collected from multiple European countries (see Huhn et al. 2014 for details). These *A. phagocytophilum* samples were previously found to represent three discreet genetic groups, which are hypothesized to have independent enzootic cycles (Huhn et al. 2014).

Prior to our analyses, from the published dataset we removed all samples that had an ambiguous nucleotide in any of the seven genetic regions, as well as the small number of samples from the United States. This left us with 278 samples in total. With these we performed a nonparametric clustering analysis using principal components to confirm sample placement into each of three discreet groups. To do this, we first concatenated the seven gene regions into a single FASTA sequence. We then used the R package Adegenet v. 2.1.3 (Jombart, 2008; Jombart and Ahmed, 2011), as implemented in R v. 4.0.2 (R Core Team, 2020), to determine the principal components from the data. The relationship between the first and second principal component was visualized in R.

### Derived strain divergence

To examine patterns of strain divergence with our dataset, we first removed any duplicated haplotype observed within each of the three strains. This left us with 10 unique haplotypes for the burrowing-mammal strain, 22 unique haplotypes for the roe deer-specialist strain, and 68 unique haplotypes for the host-generalist strain. We then performed a three-way comparison using the burrowing-mammal strain haplotypes as the outgroup to polarize observed differences. A derived site was counted if the nucleotide of the outgroup sequence at a specific position matched the nucleotide at this same position in one of the focal strain haplotypes, but the second focal haplotype had a different nucleotide. In this case, the second haplotype would be counted as a derived allele. All derived changes were classified as ‘replacement’ (i.e., they resulted in an amino acid change), or ‘silent’ (i.e., the amino acid sequence encoded by the specific codon was not different). We used a custom Perl script to make all possible 3-way comparisons among the three strains (one haplotype per strain). R v. 4.0.2 (R Core Team, 2020) was used to calculate summary statistics for the resulting output. Additionally, statistical differences in the number of observed derived sites between the two focal strains was determined with a pairwise t-test, also implemented in R v. 4.0.2 (R Core Team, 2020). Results were considered significant at p < 0.05.

### Estimation of strain divergence times

The data for *A. phagocytophilum* consisted of the same unique haplotypes as those used in our assessment of derived strain divergence (see above). To this dataset we added orthologous gene sequences from five publicly available whole genomes of the related bacterium *A. marginale* (NCBI GenBank Accessions: GCF_000020305.1, GCF_000011945.1, GCF_000495495.1, GCF_003515675.1, and GCF_003515735.1). To identify these sequences, we first located the full gene sequences for each of the seven focal genes from the annotated genome of *A. marginale* str. Florida (Accession: GCA_000020305.1). These full DNA sequences were converted to amino acid sequences and then aligned to the similarly converted, partial *A. phagocytophilum* sequences using the muscle algorithm (Edgar 2004), as implemented in SeaView v. 4.6.3 (Galtier et al. 1996, Gouy et al. 2010). The orthologous sequence regions of *A. marginale* were then used as the query sequences to locate the same regions in the other four *A. marginale* genomes with a local BLAST (‘blastp’, evalue 1e-50) (Altschul et al. 1990). With the resulting genome coordinates, we used Samtools v. 1.9 (Li et al. 2009), to extract these regions and combine them with our *A. phagocytophilum* sequences. These data were then aligned based on the amino acid sequences for the seven genetic regions as described above. Missing data and gaps indicating an insertion or deletion (INDELs) in one of the linages were removed, as were any sites that were ambiguously aligned around an INDEL, as determined by manual inspection.

Next, we determined the best partioning scheme and mutation model across the concatenated dataset using PartitionFinder 2.1.1 (Lanfear et al. 2016), with the small sample-size corrected version of the Akaike Information Criterion (AICc). The first, second, and third codon positions for the concatenated sequences were considered as separate partitions.

To estimate approximate strain splitting times, we used the Bayesian Evolutionary Analysis Utility tool (BEAUti v. 2.3.2) to create the XML file for implementation in BEAST v. 2.5.2 (Bouckaert et al., 2019). We split our dataset to apply distinct site models to the first, second and third positions. Based on the results of our PartitionFinder analysis, we used the Generalized Time Reversible (GTR) (Tavaré 1986) model of mutation for each data subset, with six gamma categories and an estimated shape. A proportion of invariant sites was also estimated for the first and second data subsets.

We ran two Markov chain Monte Carlo chains of 10^8^ iterations in the program BEAST v. 2.5.2 (Bouckaert et al., 2019), with a log normal relaxed clock and the calibrated Yule model for our tree prior. To place our divergence estimates in absolute time, we used a prior of 59 Ma (mean in real space) with a log normal distribution and an S parameter of 0.18 for the splitting time of *Anaplasma phagocytophilum* and *A. marginale*, which generated a 95% probability range that encompassed that given in Foley et al. (2008). We used LogCombiner v.1.7.5 to combine the two separate runs, and Tracer v.1.5 to ensure the effective sample size (ESS) of the parameters exceeded 200. The program TreeAnnotator v.2.5.2 was used to summarize the tree log, and FigTree v.1.4.4 (Rambaut 2018) was used for tree visualization.

### Assessment of strain-specific demographic changes

We looked at demographic changes in the two *I. ricinus*-vectored strains using an Extended Bayesian Skyline Plot (EBSP) analysis as implemented in Beast v.2.5.2 (Bouckaert et al., 2019). To do this, we used the same partitioning scheme and corresponding evolutionary models previously determined (see above). The samples that corresponded to the host-generalist and roe deer-specialist strains were examined separately. We used all samples in this analysis. To roughly estimate changes in effective population size, we applied a clock rate of 8.5 × 10^−8^ to the partition corresponding to the third codon positions of each sequence. This rate was based on a previously estimated synonymous substitution rate of 8.9 × 10^−11^ per base pair per generation determined for *Escherichia coli* (Wielgoss et al. 2011), multiplied by the 959 base pairs included in this partition. The rates for the other two partitions were estimated. A strict molecular clock rate was applied to each of the three partitions. We ran two Markov chain Monte Carlo chains of 10^8^ iterations in the program BEAST, with a burnin of 10% and sampling at every 1,000 chains. The separate runs were combined using LogCombiner v.2.5.2, and the Tracer v.1.7.1 was used to confirm that the effective sample size (ESS) of the relevant parameters exceeded 200.

### Patterns of demographic change in roe deer

To examine potential demographic changes in the European roe deer population, we utilized published sequences of the mitochondrial cytochrome b (cytb) gene amplified from 46 roe deer from Poland (Matosiuk et al. 2014). We first partitioned this dataset into the first, second and third codon sites, and used Partionfinder v.2.1.1 (Lanfear et al. 2016) to determine the most appropriate model of evolution to apply to each partition. We then used an Extended Bayesian Skyline Plot (EBSP) analysis as implemented in Beast v.2.5.2 (Bouckaert et al. 2019) to examine changes in effective population size over time in this species. The dataset was split into first, second and third positions. Following our PartionFinder results, the HKY model of substitution (Hasegawa et al. 1985), was applied to the first and second positions, and the first position additional had a portion of invariable sites (0.1) applied to it. We applied the Generalized Time Reversible (GTR) (Tavaré 1986) model of mutation to the third position, with six gamma categories and an estimated shape. To roughly estimate changes in effective population size, we applied a clock rate of 2.0 × 10^−6^ to the partition corresponding to the third codon positions of each sequence (representing 2% per site per million years) (Brown et al. 1979). We note here that use of this rough rate for mammals has been shown to be problematic (e.g., Nabholz et al. 2007), but that it is similar to comparable rates given for Artiodactyla (e.g., Bulmer et al. 1991). The rates for the other two partitions were estimated. We applied a strict molecular clock rate each of the three partitions. We ran two Markov chain Monte Carlo chains of 10^8^ iterations in the program BEAST, with a burnin of 10% and sampling at every 1,000 chains. The two separate runs were combined LogCombiner v.1.7.5, and the results assed with Tracer v.1.5 to ensure the effective sample size (ESS) of the parameters exceeded 200. We visualized the demographic changes by plotting the combined EBSP log files in R v. 4.0.2 (R Core Team, 2020), using the plotEBSP.R script which is available in the BEAST package.

## ACKNOWLEDGMENTS

N. Bates was supported in part by a FY2019 Student Faculty Scholarship Grant from Montclair State University. Q. Archer was funded by an award from the Garden State-Louis Stokes Alliances for Minority Participation (LSAMP) program (NSF Award 1909824).

